## supplemental file for "DBT is a metabolic switch for maintenance of proteostasis under proteasomal impairment"

**This file includes:**

Supplemental Figures S1 to S8
Supplemental Tables S1

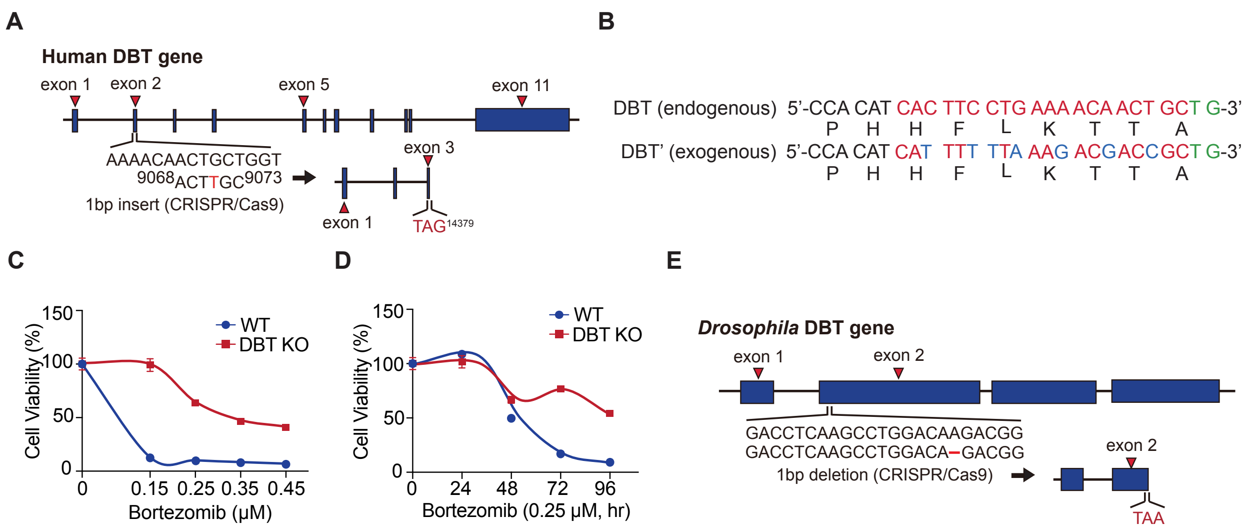


**Supplemental Figure S1. Schematics of CRISPR editing, DBT cDNA, and cell viability analysis.**

(**A**) In the DBT knockout RPE1 cell line, the human DBT gene is disrupted with a CRISPR/Cas9-induced 1-bp insert in exon 2, resulting in a premature stop codon (TAG) in exon 3 and disruption of the DBT gene in a homozygous allele. (**B**) a DBT cDNA (DBT') was engineered to harbor seven synonymous point mutations in the gRNA-targeted region, which rendered the cDNA resistant to the Cas9 cleavage in the DBT KO cell line. (**C**) The cytotoxicity analysis of WT and DBT KO RPE1 cells treated with bortezomib at different doses for 96 h. The results indicate that loss of DBT led to significant resistance to bortezomib-induced cell death (n = 3; p = 0.0386). (**D**) The cytotoxicity analysis of WT and DBT KO RPE1 cells treated with bortezomib over different periods of time (n = 3; p = 0.0182). (**E**) In the DBT knockout *Drosophila* strain, the *Drosophila* DBT gene is disrupted with a CRISPR/Cas9-induced 1-bp deletion in exon 2, resulting in a premature stop codon (TAA) in exon 2. Error bars represent ± SEM.

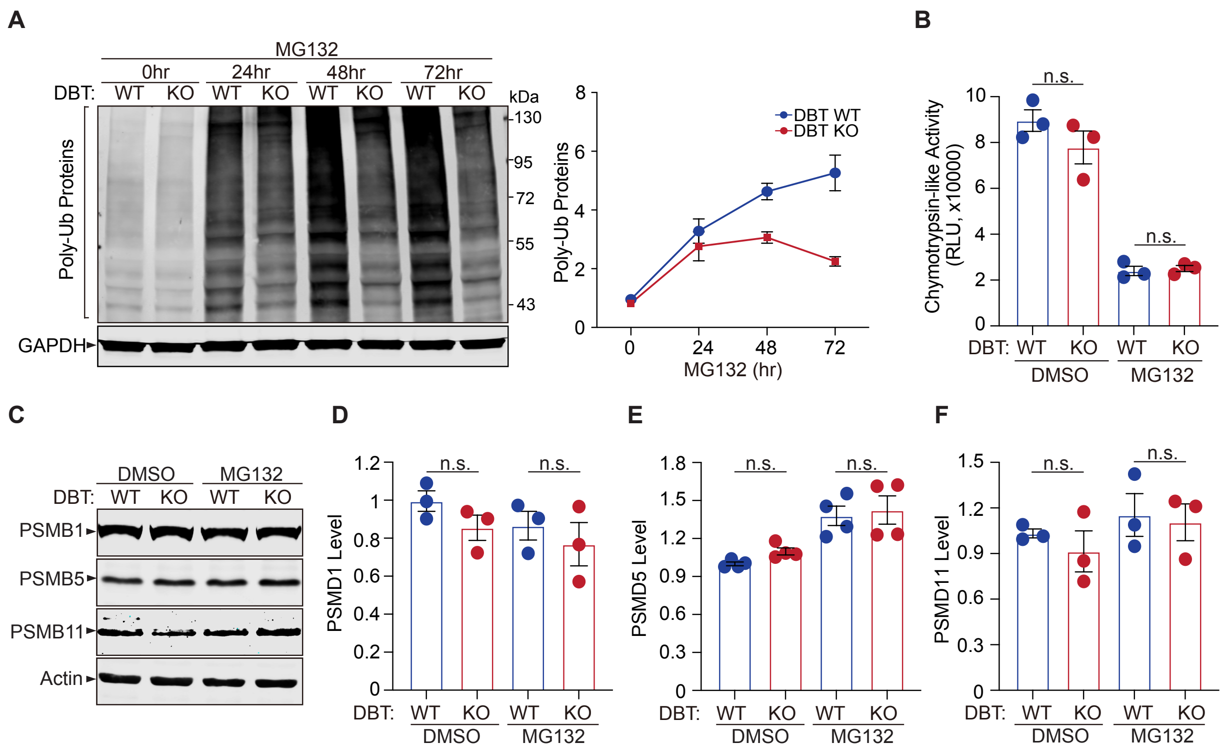


**Supplemental Figure S2. The enhanced clearance of poly-ubiquitinated proteins upon loss of DBT is not mediated by proteasomal degradation.**

(**A**) Immunoblot analysis of poly-ubiquitinated proteins in WT and DBT KO RPE1 cells treated with MG132 (2 μM) over different periods of time (n = 4; p = 0.0394). (**B**) The proteasomal activity (chymotrypsin-like) in WT and DBT KO RPE1 cells was measured with a luminogenic Proteasome-Glo substrate with or without MG132 treatment (2 μM, 48 hr) (n = 3). (**C-F**) Immunoblot analysis of proteasome subunits PSMB1, PSMB5, and PSMB11 and quantification of the subunit protein levels in the WT and DBT KO RPE1 cells treated with MG132 (2 μM, 48 hr) (n = 3-4). Error bars represent ± SEM. “n.s.”, no significance.

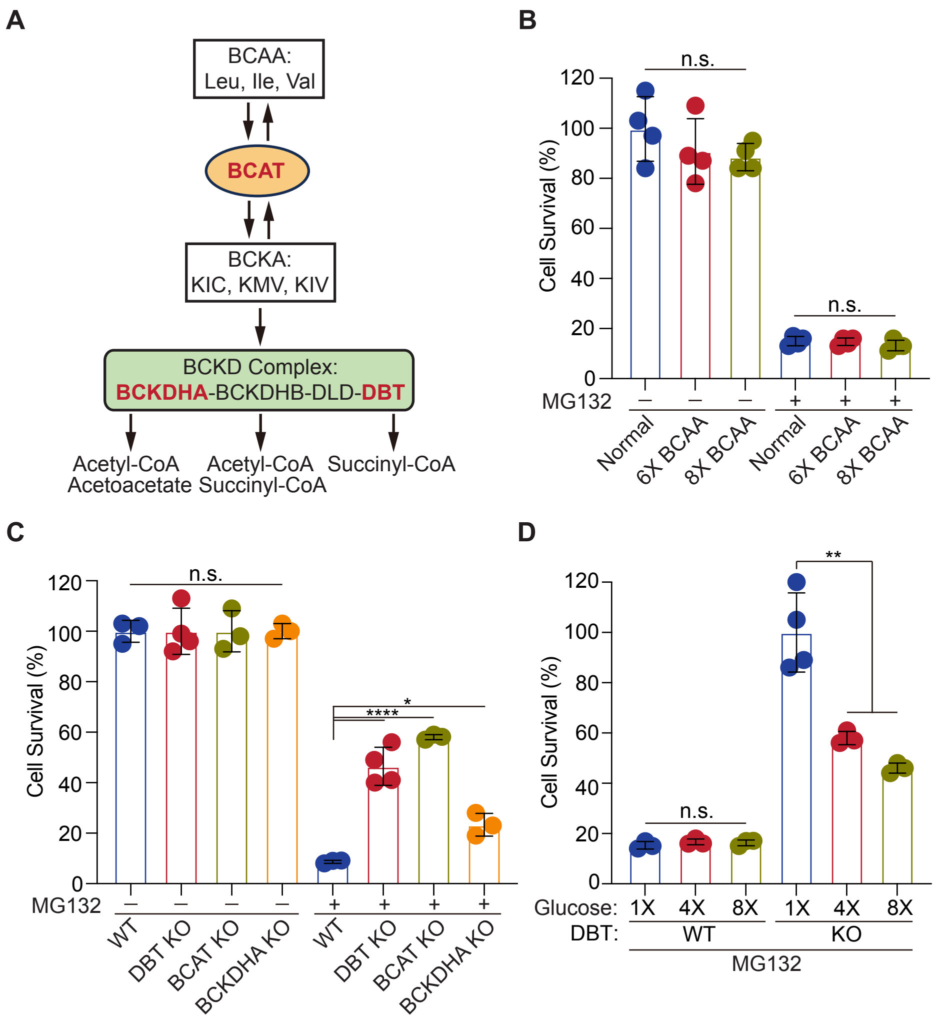


**Supplemental Figure S3. The energy depletion but not BCAA accumulation mediates the resistance of DBT knockout cells to MG132-induced proteotoxicity.**

(**A**) The pathway of BCAA catabolism: BCAAs (Leu, Ile, Val) are converted by BCAT into BCKAs (KIC, KMV, KIV), which are subsequently oxidized by the BCKD complex. Acetyl-CoA, acetoacetate, and succinyl-CoA are the final products generated through a series of subsequent reactions. (**B**) The cytotoxicity analysis of WT RPE1 cells treated with different concentrations of BCAAs (Leu, Ile, Val) (6X, 2.7 μM; 8X, 3.6 μM) with or without MG132 (2 μM) for 48 hr. The results indicate that increasing the BCAA concentration did not promote cell resistance to MG132-induced toxicity (without MG132, n = 3, p = 0.3565; with MG132, n = 3, p = 0.3758). (**C**) The cytotoxicity analysis of WT, DBT KO, BCAT KO, and BCKDHA KO RPE1 cells treated with

MG132 (2 μM) for 48 hr. The results demonstrate that DBT KO, BCAT, and BCKDHA KO cells all had increased cell resistance to MG132-induced toxicity (DBT KO, n = 4, $p < 0.0001$; BCAT KO, n = 3, $p < 0.0001$; and BCKDHA KO, n = 3, $p = 0.0125$). (**D**) The cytotoxicity analysis of WT and DBT KO cells treated with glucose at different doses (4X, 70 μM; 8X, 140 μM) with or without MG132 (2 μM) for 48 hr. These results indicate that the glucose treatment did not affect the sensitivity of WT cells to MG132-induced toxicity, but it significantly increased the sensitivity of DBT KO cells to MG132-induced toxicity. (WT, n = 3, $p = 0.467$; DBT KO, n = 3, $p = 0.0022$). Error bars represent ± SEM. “n.s.”, no significance; *$p \leq 0.05$; **$p \leq 0.01$; ****$p \leq 0.0001$.

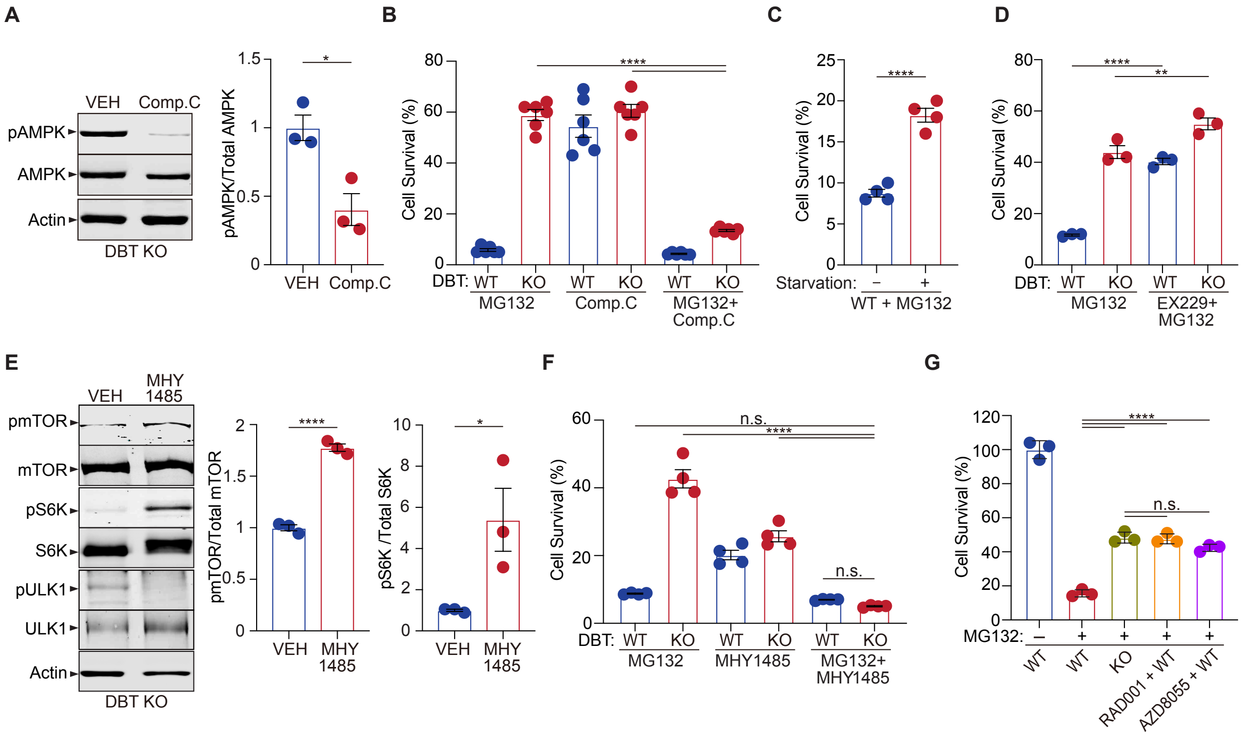


**Supplemental Figure S4. Loss of DBT activates AMPK and inhibits mTOR signaling.**

(**A**) AMPK was inactivated by the treatment with its inhibitor Compound C (10 μM, 24 hr) in DBT KO cells as indicated by the reduction in the phosphorylation of AMPK-$T^{172}$ (n = 3). (**B**) Cell viability analysis with crystal violet staining was performed on WT and DBT KO RPE1 cells treated with MG132 (2 μM, 48 hr) with or without Compound C (10 μM, 48 hr) (n = 6). (**C**) Cell viability analysis with crystal violet staining was performed on WT RPE1 cells treated with MG132 (2 μM, 48 hr) with normal medium or glucose starvation medium (n = 3). (**D**) Cell viability analysis with crystal violet staining was performed on WT and DBT KO RPE1 cells treated with MG132 (2 μM, 48 hr) with or without an AMPK agonist EX229 (10 μM, 48 hr) (n = 3). (**E**) mTOR was activated by the treatment with its activator MHY1485 (2 μM, 24 hr) in DBT KO cells as indicated by the increase in the phosphorylation of mTOR-$S^{2448}$ and its substrate S6K-$T^{389}$ (n = 3). (**F**) Cell viability analysis with crystal violet staining was performed on WT and DBT KO RPE1 cells treated with MG132 (2 μM, 48 hr) with or without MHY1485 (2 μM, 48 hr) (n = 4). (**G**) Cell viability analysis with crystal violet staining was performed on WT and DBT KO

RPE1 cells treated with MG132 (2 μM, 48 hr) with or without an mTOR inhibitor RAD001 (50 nM, 48 hr) or AZD8055 (20 nM, 48 hr) ($n = 3$). Error bars represent ± SEM. “n.s.”, no significance; *$p \leq 0.05$; **$p \leq 0.01$; ****$p \leq 0.0001$.

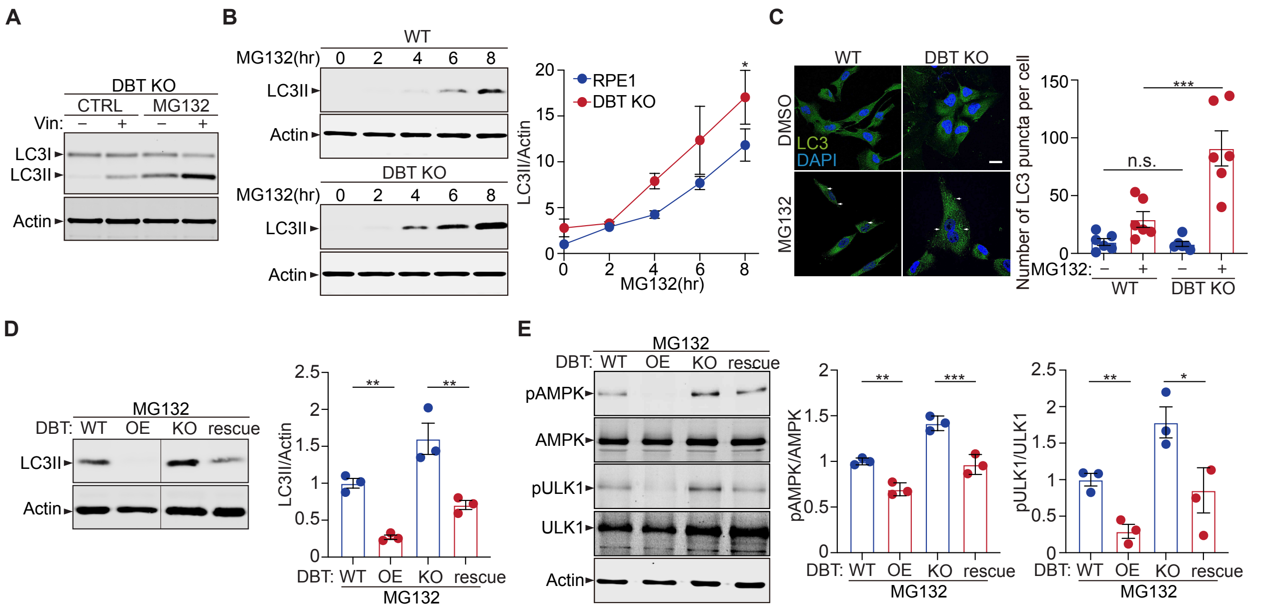


**Supplemental Figure S5. Loss of DBT promotes cellular resistance to MG132-induced toxicity through the AMPK signaling pathway.**

(**A**) Immunoblot analysis of LC3II levels in DBT KO RPE1 cells treated with MG132 (2 μM, 48 hr) with or without Vin (Vinblastine, 10 μM, 4 hr). (**B**) Immunoblot analysis of LC3II in the time-dependent analysis following MG132 treatment shows that the level of LC3II significantly increases in DBT KO cells at 4-8 hr compared to the WT control. The bar graph represents the quantification of the immunoblot analysis (n = 3). (**C**) Fluorescent immunostaining against LC3 in WT and DBT KO RPE1 cells under MG132 treatment (2 μM, 4 hr) indicates that DBT KO cells have a higher level of LC3 puncta, suggesting an increase in the autophagic function. The bar graph represents the quantification of LC3 fluorescent intensity (n = 6 experiments with >33 cells quantified for each experiment). (**D**) Immunoblot analysis of LC3II in WT and DBT KO cells transfected with DBT and co-treated with MG132 (2 μM, 48 hr) indicates that DBT overexpression (OE) in WT cells and restoration in the DBT KO cells (rescue) can reduce the LC3II levels in the WT and DBT KO cells, respectively. The bar graph represents the quantification of the immunoblot analysis (n = 3). (**E**) Immunoblot analysis of the AMPK-ULK1 axis in WT and DBT KO RPE1 cells transfected with DBT and co-treated with MG132 (2 μM, 48 hr). The activities of these regulators are quantified by measuring the levels of phosphorylation of ULK1-$S^{371}$ and AMPK-$T^{172}$ (n = 3). Scale bar: 10 μm. Error bars represent ± SEM. “n.s.”, no significance; *p ≤ 0.05; **p ≤ 0.01; ***p ≤ 0.001.

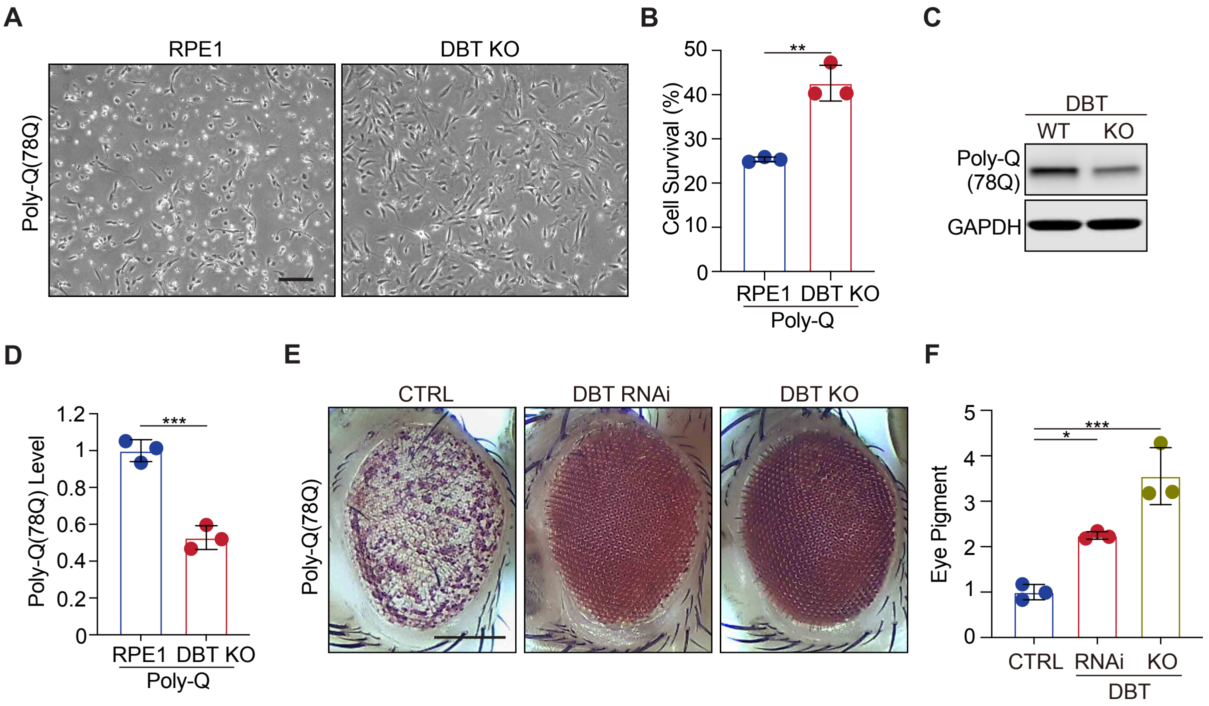


**Supplemental Figure S6. Loss of DBT protects against proteotoxicity of polyQ in mammalian neurons and *Drosophila* models.**

(**A**, **B**) Cytotoxicity analysis of polyQ expressed in WT and DBT KO RPE1 cells. The DBT KO cells exhibited higher resistance to polyQ-induced toxicity than the WT control cells ($n = 3$, $p = 0.018$). Scale bar, 300 μm. (**C**, **D**) Immunoblot analysis reveals significantly lower steady-state levels of polyQ protein in DBT KO RPE1 cells than in WT control cells. The bar graph represents the quantification of the immunoblot analysis ($n = 3$, $p = 0.0007$). (**E**, **F**) The reduction of DBT by RNAi or CRISPR led to strongly suppressed eye degeneration phenotypes in the polyQ fly strain compared to the control Luc RNAi (CTRL). The eye degeneration phenotypes were quantified by measuring the pigment content in adult eyes ($n = 3$ independent groups with each containing fly heads from 4 males and 4 females). Scale bar, 100 μm. Error bars represent means ± SEM. *$p \leq 0.05$; **$p \leq 0.01$; ***$p \leq 0.001$.

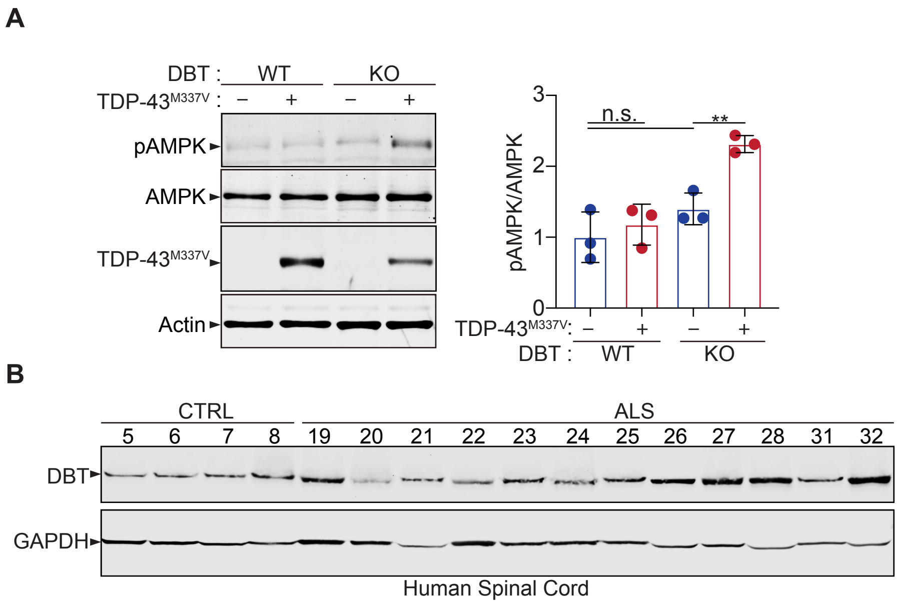


**Supplemental Figure S7. Analyses of the resistance of DBT KO cells to TDP-43 toxicity and the increased DBT protein levels in ALS patients' tissues.**

(**A**) Immunoblot analysis of the AMPK-ULK1 axis in WT and DBT KO RPE1 cells transfected with TDP-43$^{M337V}$ indicates that the DBT KO cells exhibit a higher level of phosphorylation of AMPK. The activity of AMPK is quantified by measuring the phosphorylation of AMPK-T$^{172}$ against total AMPK levels (n = 3). (**B**) Immunoblot analysis of DBT in the spinal cord tissues from ALS patients and non-neurological control cases, indicating that DBT protein levels are abnormally increased in the patient tissues. Error bars represent ± SEM. "n.s.", no significance; **p ≤ 0.01.

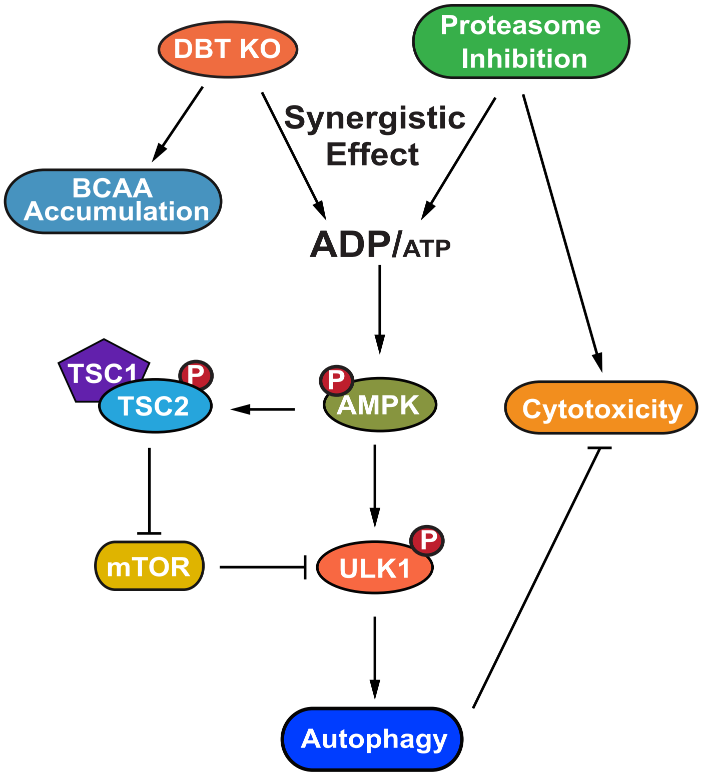


**Supplemental Figure S8. A model for the DBT-AMPK-autophagy signaling pathway.**

A working model of the mechanism through which DBT acts as a metabolic switch for the maintenance of protein homeostasis through activation of autophagy under the condition of proteasomal inhibition. The loss of DBT leads to the accumulation of BCAAs as a result of the blocked catabolism of these amino acids, which tilts the balance of intracellular energy and reduces the levels of ATP, under the condition of proteasomal inhibition. This energy imbalance triggers the activation of AMPK, which then promotes autophagy through its regulation of mTOR and ULK1. BCAA, branched-chain amino acid; ATP, adenosine triphosphate; ADP, adenosine diphosphate; AMPK, AMP-activated protein kinase; TSC1, TSC complex subunit 1; TSC2, TSC complex subunit 2; ULK1, unc-51 like autophagy activating kinase 1.

**Supplemental Table S1. The list of human patient’s tissues**.

**Supplementary Table: List of patient's tissues.**

| Sample No. | Patient ID | Source | Clinical diagnosis | ALS pathology | Age of sampling | Gender | Region |
|---|---|---|---|---|---|---|---|
| 1 | 95 | TALS | CTRL | Non | 72 | M | SC-C |
| 2 | 103 | TALS | CTRL | Non | 22 | M | SC-C |
| 3 | 110 | TALS | CTRL | Non | 50 | M | SC-C |
| 4 | 108 | TALS | CTRL | Non | 72 | M | SC-C |
| 5 | 90015 | VABBB | CTRL | Non | 66 | M | SC-C |
| 6 | 90018 | VABBB | CTRL | Non | 82 | M | SC-C |
| 7 | 100012 | VABBB | CTRL | Non | 81 | F | SC-C |
| 8 | 120016 | VABBB | CTRL | Non | 63 | F | SC-C |
| 9 | AZ160030 | VABBB | ALS | Yes (TDP-43 pathology) | 65 | M | SC-C |
| 10 | AZ140006 | VABBB | ALS | Yes (TDP-43 pathology) | 74 | M | SC-C |
| 11 | 110011 | VABBB | ALS | Yes (TDP-43 pathology) | 83 | M | SC-C |
| 12 | 140008 | VABBB | ALS | Yes (TDP-43 pathology) | 75 | M | SC-C |
| 13 | AZ150001 | VABBB | ALS | Yes (TDP-43 pathology) | 65 | M | SC-C |
| 14 | AZ140021 | VABBB | fALS | Yes | 63 | M | SC-C |
| 15 | AZ150004 | VABBB | ALS | Yes (TDP-43 pathology) | 67 | M | SC-C |
| 16 | 130022 | VABBB | ALS | Yes (TDP-43 pathology) | 48 | M | SC-C |
| 17 | 130025 | VABBB | ALS | Yes (TDP-43 pathology) | 77 | M | SC-C |
| 18 | AZ140023 | VABBB | ALS | Yes (TDP-43 pathology) | 68 | M | SC-C |
| 19 | 130020 | VABBB | ALS | Yes (TDP-43 pathology) | 78 | M | SC-C |
| 20 | 130014 | VABBB | ALS | Yes (TDP-43 pathology) | 70 | M | SC-C |
| 21 | 100007 | VABBB | fALS | Yes (TDP-43 pathology) | 61 | M | SC-C |
| 22 | 100040 | VABBB | ALS | Yes | 88 | M | SC-C |
| 23 | 120015 | VABBB | ALS | Yes (TDP-43 pathology) | 58 | M | SC-C |
| 24 | 90003 | VABBB | fALS | Yes | 73 | M | SC-C |
| 25 | 90005 | VABBB | ALS | Yes (TDP-43 pathology) | 65 | M | SC-C |
| 26 | 90020 | VABBB | fALS | Yes | 49 | F | SC-C |
| 27 | 100002 | VABBB | ALS | Yes (TDP-43 pathology) | 63 | M | SC-C |
| 28 | AZ140017 | VABBB | ALS | Yes (TDP-43 pathology) | 66 | M | SC-C |
| 29 | 38 | TALS | sALS | Yes (C9orf72 HRE) | 34 | F | SC-C |
| 30 | 88 | TALS | sALS | Yes (C9orf72 HRE) | 59 | M | SC-C |
| 31 | 92 | TALS | fALS | Yes (C9orf72 HRE) | 72 | M | SC-C |
| 32 | MY9 | TALS | ALS | Yes (C9orf72 HRE) | 62 | F | SC-C |

Abbreviations: ALS (Amyotrophic Lateral Sclerosis, fALS (familial amyotrophic lateral sclerosis), sALS (sporadic amyotrophic lateral sclerosis), CTRL (Control), M (Male), F (Female), TALS (Target ALS Human Postmortem Tissue Core), VABBB (VA Biorepository Brain Bank), SC-C (Spinal Cord-Cervical), and HRE (hexanucleotide repeat expansion). The mean age of the controls is 63.5 years versus 69.4 years for the patients.
